# Scalable and cost-effective ribonuclease-based rRNA depletion for transcriptomics

**DOI:** 10.1101/645895

**Authors:** Yiming Huang, Ravi U Sheth, Andrew Kaufman, Harris H Wang

## Abstract

Bacterial RNA sequencing (RNA-seq) is a powerful approach for quantitatively delineating the global transcriptional profiles of microbes in order to gain deeper understanding of their physiology and function. Cost-effective bacterial RNA-seq requires efficient physical removal of ribosomal RNA (rRNA), which otherwise dominates transcriptomic reads. However, current methods to effectively deplete rRNA of diverse non-model bacterial species are lacking. Here, we describe a probe and ribonuclease based strategy for bacterial rRNA removal. We implemented the method using either chemically synthesized oligonucleotides or amplicon-based single-stranded DNA probes and validated the technique on three novel gut microbiota isolates from three distinct phyla. We further showed that different probe sets can be used on closely related species. We provide a detailed methods protocol, probe sets for >5,000 common microbes from RefSeq, and an online tool to generate custom probe libraries. This approach lays the groundwork for large-scale and cost-effective bacterial transcriptomics studies.

## Introduction

Bacterial RNA-seq provides global and in-depth transcriptional profiling of a given species or community that can yield quantitative and mechanistic insights into microbial function and ecology, and holds exciting potential for routine use in microbiology studies and large-scale screening applications(1–7). In bacteria, the 16S and 23S ribosomal RNA (rRNA) are the most abundant RNA molecules in the cell and typically account for more than 90% of the total RNA(8). These rRNA therefore need to be experimentally removed from total RNA during library preparation steps to ensure cost-effective bacterial RNA-seq with sufficient coverage of coding sequences of the transcriptome. Mammalian RNA-seq has a similar challenge of rRNA removal, but solutions are more tractable since 1) most mammalian mRNAs are polyadenylated and can thus be easily selected or captured(9,10), or 2) the same species (e.g. human, with the same rRNA sequences) are generally profiled such that sequence-specific depletion methods can be streamlined(11–14). In contrast, bacterial mRNAs do not have a poly(A) sequence, which makes enrichment strategies infeasible, and 16S and 23S rRNA sequences are diverse between species, which makes universal depletion approaches much more challenging. A number of commercially available rRNA depletion methods exist(8,15–18) and several different depletion strategies have been developed (Table 1). MICROBExpress (Ambion), RiboMinus (Life Technologies) and riboPOOL (siTOOLs Biotech) utilize probes targeting 16S and 23S rRNA to capture rRNA; mRNA-ONLY (Epicentre) uses a 5’-monophosphate-dependent exonuclease to degrade bacterial rRNA molecules; and Ovation Prokaryotic RNA-seq System (NuGEN) uses selective primer sets to avoid targeting rRNA in cDNA synthesis. These kits can be used to deplete bacterial rRNA for single-species RNA-seq and environmental metatranscriptome profiling, and a subset can be used with custom designed probes to target non-model species(19–22).

**Table 1:** rRNA depletion methodologies for bacterial transcriptomics

| Name | Catalog | Depletion strategy | Compatibility (fragmented RNA) | Cost per reaction | Notes |
| --- | --- | --- | --- | --- | --- |
| This work | - | RNase H degradation | Yes | ~\$13 | costs ~\$5 more for amplicon probes |
| Ribo-Zero™ rRNA Removal Kit (Bacteria) | - | Subtractive hybridization | Yes | ~\$80 | recently discontinued |
| MICROBExpress Bacterial mRNA Enrichment kit | ThermoFisher AM1905 | Subtractive hybridization | No | ~\$24 | targets short conserved regions of rRNA |
| mRNA-ONLY Prokaryotic mRNA Isolation kit | Epicentre MOP51010 | Exonuclease digestion | No | Inquiry required | based on 5'-monophosphate-dependent rRNA degradation |
| Trimmer-Direct cDNA Normalization Kit | Evrogen NK003 | DSN-based normalization(24) | Yes | ~\$33 | potentially GC biased and low-efficiency(8) |
| Ovation Prokaryotic RNA-seq System | NuGEN 0326-32 | Selective cDNA synthesis | No | ~\$95 | price includes reagents for library preparation |
| RiboMinus™ Transcriptome Isolation Kit, bacteria | ThermoFisher K155004 | Subtractive hybridization | No | ~\$48 | targets short conserved regions of rRNA |
| Pan-Prokaryote riboPOOL | siTOOLS Biotech | Subtractive hybridization | No | ~\$13 | targets partial regions of rRNA |

However, all of these techniques rely on targeting short conserved rRNA regions or specific rRNA modifications such that only full length rRNA can be removed. Recent methods for RNA barcoding have enabled dramatically reduced cost in RNA-seq by pooling multiple samples together at the total RNA stage and performing downstream library preparation reactions in a pooled format(23). However, since these protocols generate fragmented RNA (including rRNA), these standard rRNA removal kits cannot be used. The Ribo-Zero kit (Illumina, Inc.) was a popular commercial rRNA depletion kit that was compatible with fragmented RNA and could be used effectively on some (e.g. (23)) but not all bacteria. Unfortunately, this kit was recently discontinued by the manufacturer. While sequence-independent rRNA depletion methods such as those that rely on DSN nuclease have been described(24), they suffer from low efficiency and sequence-specific or GC biases, e.g. the DSN method exhibits poor rRNA depletion efficiency (> 75% rRNA reads after depletion) and severe GC biases of transcript enrichment (R2 = 0.13) for GC-rich *R. sphaeroides*(8).

Motivated by the lack of a reliable and low-cost alternatives, we set out to develop a bacterial rRNA depletion method with the following criteria: 1) high efficiency 16S/23S rRNA depletion, 2) scalable to diverse bacterial species, 3) compatibility with fragmented or barcoded RNA samples, and 4) low cost per reaction. Correspondingly, we developed a ribonuclease (RNase) H based method for bacterial rRNA depletion described here. We show that our approach efficiently depletes bacterial rRNA with minimal off-target effects to the transcriptome and demonstrate the use of the method on bacterial species from three different phyla. Moreover, we show that species-specific depletion probes can be generated by chemical synthesis or from PCR amplicons (with different cost considerations) and that probe-sets can be utilized on species with similar 16S/23S sequences.

## Materials and Methods

### Bacterial strains and culture condition

*Bacteroides dorei*, *Collinsella aerofaciens* and *Dorea longicatena* are novel isolates derived from a fecal sample obtained from a healthy human adult subject, obtained during the course of unrelated experiments. This work was approved and conducted under Columbia University Medical Center Institutional Review Board protocol AAAR0753, and written informed consent was obtained from the subject. *Bacteroides uniformis* strain ATCC 8492 and *Bacteroides vulgatus* strain ATCC 8482 were obtained from American Type Culture Collection, Manassas, VA, USA. All bacterial strains were grown in Gifu Anaerobic Medium Broth, Modified [HyServe 05433] under anaerobic conditions (5% H2, 10% CO2, 85% N2) in a Coy Laboratory Products anaerobic chamber. To characterize the *B. dorei* strain under different perturbations, cells were cultured in Gifu Anaerobic Medium Broth, Modified [HyServe 05433] under anaerobic conditions (5% H2, 10% CO2, 85% N2), and short-chain fatty acids or carbohydrates were added to cultures at a final concentration of 5 mg/mL at exponential phase. Cell cultures were harvested for RNA extraction after 90 minutes of exposure and immediately frozen at −80°C.

### Bacterial RNA and DNA extraction

Bacterial RNA was obtained utilizing the RNAsnap method(25). Briefly, 1mL of saturated culture was centrifuged and the cell pellet was stored at −80°C. The pellet was then suspended in 500μL RNAsnap mix (95% formamide, 18mM EDTA, 0.025% SDS and 1% B-mercaptoethanol), lysed using ~200μL of 0.1mm Zirconia Silica beads [Biospec 11079101Z] using a BioSpec mini bead beater [Biospec 1001] for 2 minutes in a deep well 96-well plate [Axygen P-DW-20-C]. The plate was centrifuged and the clarified lysate was cleaned up using a Zymo ZR-96 RNA Clean & Concentrator kit [Zymo R1080] per manufacturer’s instructions. Bacterial DNA was obtained using our standard automated DNA extraction pipeline based on a scaled down version of the Qiagen MagAttract PowerMicrobiome DNA/RNA Kit [Qiagen 27500-4-EP], detailed fully in (26).

### Genome sequencing and assembly of novel isolates

To obtain draft genome references for novel isolates, paired-end libraries were constructed using a low-volume Nextera sequencing protocol(27) and sequenced on Illumina Nextseq 500/550 platform to obtain about 6 million paired-end reads with a read length of 2×75bp for each novel isolate. The reads were processed using Cutadapt v2.1(28) with following parameters “-a file:[Nextera_adapter.fa] -A file:[Nextera_adapter.fa] --minimum-length 25:25 -u 15 -u −5 -U 15 -U −5 -q 15 --max-n 0 --pair-filter=any” to remove first 15 bases, last 5 bases and adapters. Nextera adapter sequences used for trimming are provided in the Supplementary Table 4. To further improve the performance of genome assembling, PacBio long-read sequencing was performed for each isolate by SNPsaurus. PacBio long reads and remaining Illumina reads were assembled using Unicycler(29) in hybrid mode with default parameters to generate draft assembly for each isolate. The draft assemblies were further assessed using CheckM(30) to ensure their high quality (completeness >99% and contamination <1%).

### Draft genome assembly annotation

Draft genomes for novel isolates (*Bacteroides dorei*, *Collinsella aerofaciens* and *Dorea longicatena)* were *de novo* assembled as described above, and genome assemblies for *Bacteroides uniformis* strain ATCC 8492 and *Bacteroides vulgatus* strain ATCC 8482 were downloaded from NCBI (RefSeq assembly accession: GCF_000154205.1 and GCF_000012825.1). Genome annotation was performed using Prokka v1.13.3(31) with following parameters “--rnammer --rfam” for all 5 bacterial strains, and sequences of predicted 16S and 23S rRNAs were extracted from genome using in-house script for downstream rRNA reads alignment (only one 16S and one 23S rRNA were used for strains with multiple rRNA annotations since their sequences are almost identical). Polysaccharide-utilization loci (PUL) and susC/D gene pairs in *B. dorei* were predicted by PULpy(32).

### RNA-seq library preparation and sequencing

The RNA-seq library was constructed following standard RNAtag-seq protocol with minor modifications, detailed fully in (23).

Briefly, 4μg of total RNA was fragmented using 2X FastAP buffer [ThermoFisher EF0651], depleted of DNA and dephosphorylated using TURBO™ DNase [ThermoFisher AM2239] and FastAP [ThermoFisher EF0651], and then fragmented RNA after 2X SeraPure SPRI beads cleanup(33,34) was ligated to barcoded first adapter [NEB M0437M], yielding barcoded RNA. Uniquely barcoded RNA from different conditions were pooled together and further subjected to size selection and purification using the Zymo RNA Clean and Concentrator-5 [Zymo R1015]. Pooled, barcoded RNA was then quantified using Qubit RNA HS Assay Kit [ThermoFisher Q32855] per the manufacturer’s instructions.

Normalized barcoded RNA samples were previously subjected to rRNA depletion using Illumina Ribo-Zero rRNA Removal Kit (Bacterial) per the manufacturer’s instructions in standard RNAtag-seq protocol(23), and we replaced the typical Ribo-Zero rRNA depletion step with our RNase H depletion reaction. The same starting RNA samples on which Ribo-Zero depletion was previously performed were used for the RNase H reaction, and 100ng of barcoded RNA was subjected to RNase H based rRNA depletion for condition optimization experiment while 500ng of barcoded RNA was subjected to RNase H based rRNA depletion to validate depletion performance across diverse bacterial species and with different types of probes. To generate library with no rRNA depletion as control, 16ng of barcoded RNA (for optimization experiment) and 50ng RNA of barcoded RNA (for validation experiment) was directly diluted into the final volume and subjected to downstream library preparation. For the *B. dorei* perturbation experiment, 300ng of pooled, barcoded RNA was subjected to RNase H based rRNA depletion.

rRNA depleted RNA was subjected to downstream library preparation following standard RNAtag-seq protocol(23), including reverse transcription [ThermoFisher 18090010], degradation of the RNA [NEB M0297S] and ligation of second adapter [NEB M0437M]. Ligation product was further amplified with primers that contain Illumina P5 and P7 adapters and sample indexes, and the PCR reactions were stopped during exponential amplification (typically ~15 cycles). The PCR products were subjected to gel electrophoresis on E-Gel™ EX Agarose Gels, 2% [ThermoFisher G402002] and expected DNA bands (around 300bp-600bp) were excised from gel and extracted by Wizard™ SV Gel and PCR Cleanup System [Promega A9282] following the manufacturer’s instructions to remove PCR primers and adapter dimers from sequencing library. Resulting libraries were sequenced to a low coverage in optimization experiments and a high coverage in validation experiments and screening experiments (Supplementary Table 5). Sequences of all adapters and primers used in library preparation are provided in the Supplementary Table 6; we note that in our implementation of RNAtag-seq we have swapped the orientation of R1 and R2 primers such that RNA barcodes are sequenced in read 2 and not read 1.

For the condition optimization experiment (100ng of uniquely barcoded RNA as input for RNase H reaction) except the un-depleted control, libraries were sequenced on Illumina MiSeq platform (reagent kits: v2 50-cycles, single-end mode) at 12 pM loading concentration with 10% PhiX spike-in (PhiX Control V3 [Illumina FC-110-3001]) following the manufacturer’s instructions.

For the depletion performance validation experiment (500ng of uniquely barcoded RNA as input for RNase H reaction) and the un-depleted control in optimization experiment, libraries were sequenced on Illumina NextSeq 500/550 platform (reagent kits: High Output v2 75-cycles, single-end mode) at 1.8 pM loading concentration with 2.5% PhiX spike-in (PhiX Control V3 [Illumina FC-110-3001]) following the manufacturer’s instructions.

For the *B. dorei* screening experiment (300ng of pooled barcoded RNA as input for RNase H reaction), libraries were sequenced on Illumina NextSeq 500/550 platform (reagent kits: High Output v2 75-cycles, paired-end mode with R1 67 and R2 8) at 1.8 pM loading concentration with 2.5% PhiX spike-in (PhiX Control V3 [Illumina FC-110-3001]) following the manufacturer’s instructions. Paired-end reads were further demultiplexed based on the barcode sequences in R2 using sabre(35) with at most one mismatch allowed in the barcodes.

### ssDNA probe generation

Probes for a species of interest can be generated via two strategies: chemically synthesized oligonucleotide (oligo) or amplicon-based ssDNA (amplicon).

For oligo probes, we developed a tool to automatically design oligo probe libraries for given 16S and 23S sequences: ribosomal RNA sequence was spilt into multiple ~50nt DNA oligos covering the entire length of its reverse complement. The oligo probe libraries for 5 strains were chemically synthesized from Integrated DNA Technologies Inc. in plate format with normal desalting purification, and equimolarly pooled together to generate oligo probe mixes used in this study.

For amplicon probes, universal primers in which the forward primer is 5’-phosphorylated (8F/1541R for 16S amplicon and 10F/2756R for 23S amplicon(36), Supplementary Table 3) were used to amplify 16S and 23S amplicons from bacterial genomic DNA, and the resulting PCR product was subjected to gel electrophoresis (E-Gel™ EX Agarose Gels, 2% [ThermoFisher G402002]), and gel extraction (Wizard™ SV Gel and PCR Cleanup System [Promega A9282]) following the manufacturer’s instructions, then 16S and 23S amplicons were amplified again using the same primers with gel extract as template. Amplicon products were purified using 1X SeraPure SPRI beads(33,34) and then subjected to a lambda exonuclease [ThermoFisher EN0562] digestion, which selectively digests the 5’-phosphorylated strands resulting in amplicon ssDNA probes complementary to ribosomal RNAs. Finally, 16S amplicon probe and 23S amplicon probe were mixed in equimolar amounts to generate amplicon probe mixes used in this study. A detailed step-by-step protocol for the method is included in Supplementary Methods.

### RNase H reaction optimization

To optimize RNase H reaction conditions, we added oligo probe mix to 100ng of fragmented RNA after first adapter ligation with various probe-to-RNA ratios (1:1, 5:1 or 10:1), the mixture was incubated in hybridization buffer (200mM NaCl, 100mM Tris-HCl pH7.5) in a final volume of 5μL at 95°C for 2 minutes, the temperature was slowly ramped down (−0.1°C/sec) to 45°C (for Hybridase RNase H) or 22°C (for NEB RNase H), and the mixture was incubated at 45°C (for Hybridase RNase H) or 22°C (for NEB RNase H) for additional 5 minutes. After probe hybridization, we added 5μL preheated Hybridase RNase H reaction mix (10U Hybridase Thermostable RNase H [Lucigen H39500], 0.5μmol Tris-HCl pH 7.5, 1μmol NaCl and 0.2μmol MgCl2) or 5μL NEB RNase H reaction mix (10U NEB RNase H [NEB M0297S] and 1uL 10X NEB RNase H reaction buffer) to the RNA-probe mixture and incubated the mixture at 45°C (for Hybridase RNase H) or 37°C (for NEB RNase H) for various depletion times (10 minutes, 30 minutes and 60 minutes). After RNase H digestion, 15μL DNase I reaction mix (4U TURBO DNase [ThermoFisher AM2239] and 2.5μL 10X TURBO DNase Buffer) was added, and the mixture was incubated at 37°C for 30 minutes to degrade ssDNA probes. RNA after depletion was purified with 2X SeraPure SPRI beads(33,34) and eluted into 14μL for downstream library construction.

### RNase H based ribosomal RNA depletion

Based on the results of optimization experiments, we selected reaction conditions of Hybridase RNase H enzyme, 5:1 probe-to-RNA ratio and 30 minutes reaction duration. To validate the performance of this method across diverse bacterial species and with different types of probes, we used 500ng of RNA as input and scaled the hybridization reaction to 15μL. We added 2500ng ssDNA probe mix (oligo or amplicon) to 500ng of fragmented RNA after first adapter ligation, the mixture was incubated in hybridization buffer (200mM NaCl, 100mM Tris-HCl pH7.5) in a final volume of 15μL at 95°C for 2 minutes, the temperature was slowly ramped down (−0.1°C/sec) to 45°C, and the mixture was incubated at 45°C for additional 5 minutes. After probe hybridization, we added 5μL preheated Hybridase RNase H reaction mix that contains 15U Hybridase Thermostable RNase H [Lucigen H39500], 0.5μmol Tris-HCl pH 7.5, 1μmol NaCl and 0.4μmol MgCl2 to the RNA-probe mixture and incubated the mixture at 45°C for 30 minutes. After RNase H digestion, 30μL DNase I reaction mix (6U TURBO DNase [ThermoFisher AM2239] and 5μL 10X TURBO DNase Buffer) was added, and the mixture was incubated at 37°C for 30 minutes to degrade ssDNA probes. RNA after depletion was purified with 2X SeraPure SPRI beads(33,34) and eluted into 14μL for downstream library construction. For RNA ligated to DNA adapters, DNase I digestion is replaced by 2X SeraPure SPRI beads cleanup to remove oligo probes (e.g. to avoid degrading the DNA adapter barcode). A detailed step-by-step protocol for the method is included in Supplementary Methods.

### RNA-seq data analysis

Raw sequencing reads were first processed using Cutadapt v2.1(28) with following parameters “-a file:[RNAtagSeq_adapter.fa] -u 5 --minimum-length 20 --max-n 0 -q 20” to remove first 5 bases and adapters before alignment. RNAtag-seq adapter sequences used for trimming are provided in the Supplementary Table 4. To remove reads originated from PhiX spike-in, we aligned remaining reads to the Coliphage phi-X174 genome (downloaded from NCBI, sequence accession: NC_001422.1) using Bowtie2 v2.3.4 in single-end and “--very-sensitive” mode, and only unmapped reads were used for downstream alignment. We then aligned remaining reads to 16S and 23S rRNAs of corresponding strains using Bowtie2 v2.3.4(37) in single-end mode to estimate the proportion of reads originating from rRNA. The reads not mapped to rRNA were also outputted by “--un” option in Bowtie2 and further aligned to corresponding genome reference using Bowtie v1.2.2(38) in single-end mode with following parameters “-a -m 1 --best --strata”. All alignments were subsequently converted to BAM format, sorted and indexed using SAMtools v1.9(39) for further analysis.

To evaluate the performance of rRNA depletion, following read-level metrics were calculated (reads with multiple alignments suppressed in BAM file were included in the following calculations): proportion of reads originating from rRNA, i.e. rRNA reads (%), was defined as: number of reads mapped to rRNA / number of total mapped reads; and fold enrichment of non-rRNA reads was defined as: fold change of (1 - proportion of reads originating from rRNA).

To evaluate the transcriptomic consistency between depleted samples and un-depleted samples, the number of reads uniquely mapped to each CDS (reads with multiple alignments were excluded) was calculated using featureCounts v1.6.2(40) without restraint on strandness (− s 0), we then quantified the expression level of each CDS by transcripts per million (TPM) using in-house script. We used log10(TPM + 0.1) as input to calculate the Pearson correlation, Spearman correlation and R-squared for gene-level correlation measurement, and visualized the quantiles of TPM distributions in quantile-quantile (Q-Q) plots to evaluate whether rRNA depletion could cause global shift in transcriptome. Effect sizes of Mann-Whitney U tests performed on depleted and un-depleted samples were also calculated to quantify the distribution shift in addition to Q-Q plots(41).

To characterize the impact of short-chain fatty acids and carbohydrates on *B. dorei* gene expression, we used TPM profiles as input to calculate the pairwise Spearman correlation s between conditions and performed multidimensional scaling on normalized Spearman correlation 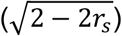 to visualize the overall effect of substrates on bacterial transcriptome. We then used DESeq2(42) to perform differential expression analysis on the number of reads uniquely mapped to each CDS, and calculated the fold-change of gene expression after perturbations. P-values of differential expression were adjusted using Benjamini-Hochberg procedure in DESeq2 using default settings.

### Probe-to-target sequence similarity analysis

To test the scalability of probes to other species, we quantified probe-to-target sequence similarity by the number of mismatches in their sequence alignment. Firstly, 16S and 23S rRNA sequences of *B. dorei* and target species were aligned respectively using MUSCLE v3.8.31(43) with default setting, then we counted the number of mismatches (including gaps) in their alignment, and the percent of mismatches in rRNAs was calculated as: number of mismatches in rRNA alignment / total length of rRNA alignment (including gaps). In addition, for each individual probe in *B. dorei* oligo library, we counted the number of mismatches in the ~50nt tile of rRNA alignment that this probe covers (including gaps), and percent of mismatches in each probe was calculated as: the number of mismatches in the tile (including gaps) / total length of the tile (including gaps). In addition, depletion efficiency, i.e. rRNA fold change, of that local rRNA sequence was calculated as the average fold change of reads counts (normalized to non-rRNA mapped reads) for each base in the tile.

### Probe specificity and off-targets analysis

To predict potential off-target depletion, we mapped the probe sequences to bacterial transcriptome generated by Prokka v1.13.3(31) using two aligners: 1) BLASTN 2.9.0+(44) with default settings and 2) BURST v0.99.8 DB15(45) with the following settings “-fr -i 0.80 -m FORAGE”. Transcripts to which probe sequences are mapped by either aligner with no more than 8 mismatches are considered as potential off-targets.

## Results

### A RNase H based rRNA depletion strategy

RNase H based methods are widely used for mammalian rRNA depletion(11,13). We thus aimed to implement and optimize a similar strategy for bacterial rRNA depletion. Given the diversity of bacterial 16S and 23S sequences, our method requires a user-customizable approach to design new probe sequences. Probes for a species of interest can be generated via two strategies (Fig. 1a). First, in a chemical synthesis approach, probes can be designed from 16S and 23S sequences by splitting sequences into ~50nt single-stranded (ss) DNA oligonucleotides that are synthesized de novo. Second, in a PCR amplicon-based approach, universal primers (the forward primer is 5’-phosphorylated) can be used to amplify 16S and 23S amplicons from genomic DNA and the resulting product can be converted to ssDNA by lambda exonuclease, which selectively digests 5’-phosphorylated strands of dsDNA(46). The generated ssDNA probes (oligo or amplicon) are subsequently hybridized to total RNA. The ssDNA:ssRNA duplexes are selectively degraded with RNase H, and the remaining ssDNA probes in the reaction are then degraded by DNase I, thus finally yielding enriched mRNAs (Fig. 1b).

**Figure 1:**
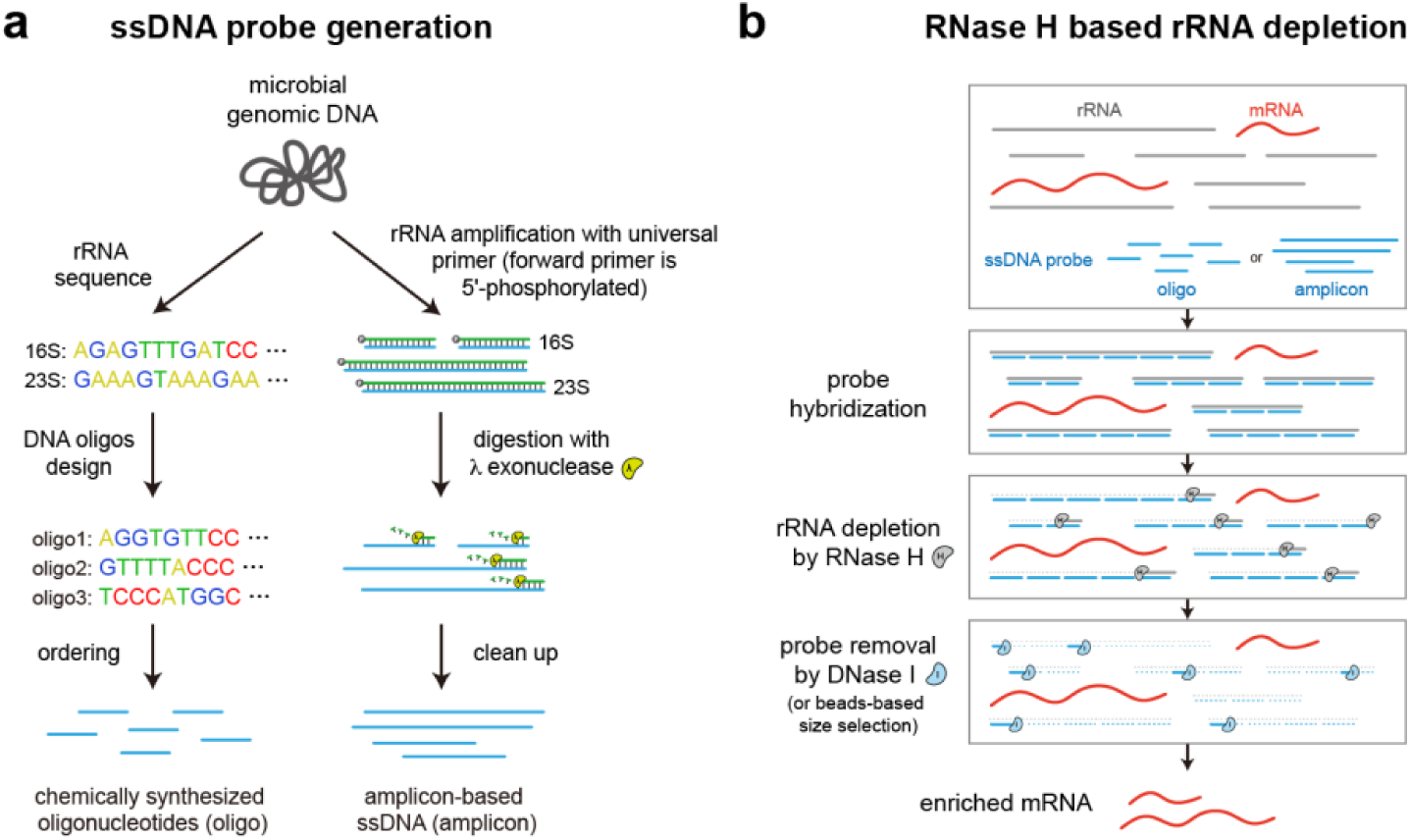
Workflow for bacterial RNase H based rRNA depletion. **a)** Probes used for depletion can be either designed and chemically synthesized from known rRNA sequences (oligo-based) or generated by PCR from genomic DNA with 5’-phosphorylated forward primers and subsequent lambda exonuclease digestion (amplicon-based). **b)** Probes are then hybridized to total RNA and the rRNA bound by the ssDNA probes is degraded by RNase H. Finally, all remaining probes are degraded by DNase I or removed by SPRI beads-based size selection, resulting in enriched mRNAs.

### Optimizing RNase H reaction parameters

We first optimized the conditions for probe hybridization and RNase H degradation of bacterial rRNA. To test reaction conditions, we took total RNA isolated from a novel human gut microbial isolate, *Bacteroides dorei*, and split the same RNA across multiple conditions. 16S and 23S sequences were determined from a draft genome of the isolate and a probe library composed of 87 distinct ~50nt oligonucleotides that tiled the 16S and 23S sequences was designed and chemically synthesized (Methods, Supplementary Table 1). We chose to use ~50nt probes as they have shown favorable performance in previous mammalian rRNA depletion strategies(13). Total RNA from this strain was subjected to fragmentation and adapter ligation (with sample barcoding) as per the standard RNAtag-seq protocol (Methods)(23). We replaced the typical Ribo-Zero rRNA depletion step with our RNase H depletion reaction, and the resulting RNA after depletion was subjected to reverse transcription, second adapter ligation, PCR amplification, and Illumina sequencing (to a low coverage for initial optimization, Methods). The resulting sequencing reads were analyzed using a standard RNA-seq analysis pipeline and the proportion of reads originating from rRNA of total mapped reads was determined (Methods).

We tested two different RNase H enzymes, a traditional RNase H (New England Biolabs) and a thermostable HybridaseTM RNase H (Lucigen), with their corresponding buffer and temperature conditions at a fixed probe-to-RNA of 1:1 based on previous mammalian RNase H depletion protocols(13) (Methods). In addition, we tested various reaction times (Fig. 2a). For both RNase H enzymes, the proportion of rRNA reads decreased after depletion, with longer incubation times yielding improved depletion efficiency for Hybridase RNase H but not NEB RNase H. To understand the contribution of probe-to-RNA ratio to overall depletion efficiency, we fixed the reaction time to 30 minutes and tested different probe-to-RNA ratios (Fig. 2b). For both RNase H enzymes, increasing probe-to-RNA ratios improved depletion efficiency, yielding low levels of reads mapped to rRNA (NEB 5:1 probe: 12.90%, Hybridase 5:1 probe: 6.82%); overall the Hybridase RNase H enzyme outperformed the NEB RNase H enzyme at all ratios. Based on these results, we adopted the following depletion enzyme and conditions for subsequent studies: Hybridase RNase H with its corresponding reaction buffers, a probe-to-RNA ratio of 5:1, and a reaction duration of 30 minutes. While even higher probe-to-RNA ratios or longer reaction times led to better performance, we chose to use the minimal conditions need to yield satisfactory depletion in order to minimize potential off-target degradation.

**Figure 2:**
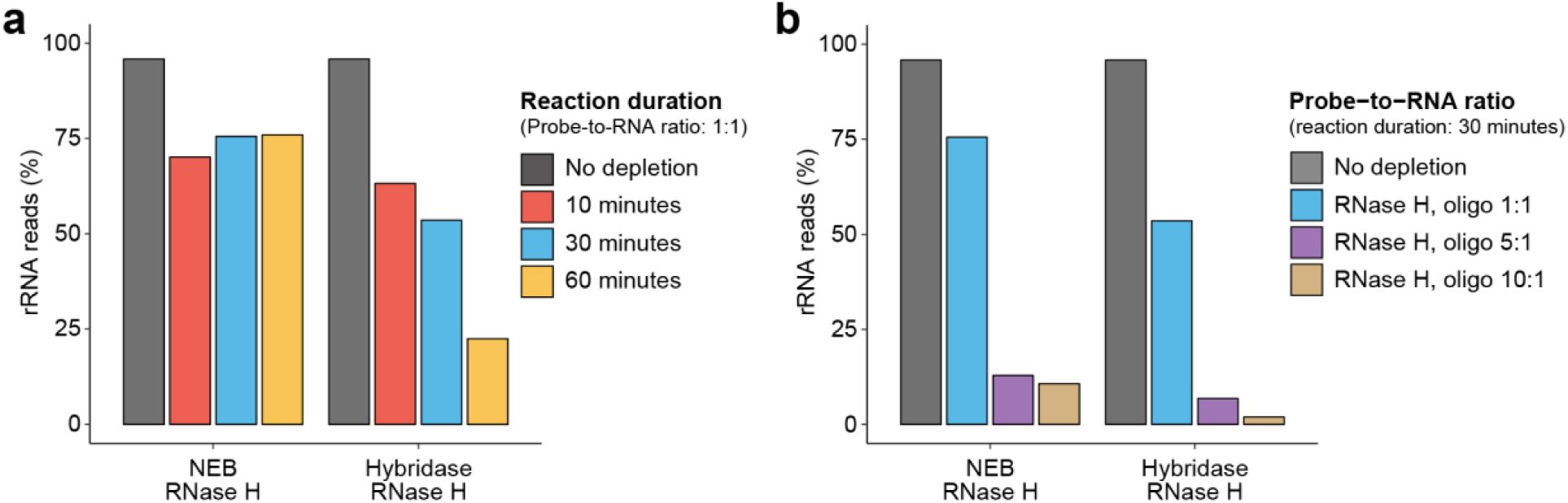
Optimization of RNase H reaction conditions. **a)** Proportion of rRNA-aligning reads of total mapped reads (% rRNA reads), for un-depleted and RNase H depleted samples for two RNase H enzymes and various reaction times with probe-to-RNA fixed to 1:1, or **b)** various probe-to-RNA ratios with reaction time fixed to 30 minutes. The same RNA sample isolated from *Bacteroides dorei* was split for rRNA depletion across different reaction conditions.

### Validating RNase H-based rRNA depletion

Having determined an optimal set of reaction conditions, we next set out to validate the rRNA depletion performance of the method across diverse bacterial species and assess any potential technical artifacts the method may introduce. We obtained a new RNA sample for the *B. dorei* strain, and also obtained RNA from two additional distinct novel gut microbiota isolates for testing, *Collinsella aerofaciens* and *Dorea longicatena.* Collectively these three isolates represent three distinct phyla (Bacteroidetes, Actinobacteria, Firmicutes, respectively). We performed library preparation and rRNA depletion as before, utilizing the optimal reaction conditions as well as a chemically synthesized oligonucleotide probe library designed for each strain based on draft genomes from Illumina assemblies (Supplementary Table 1). The resulting RNA libraries as well as the same RNA with no rRNA depletion were sequenced at a high coverage (Methods). In all three strains, our method significantly depleted rRNA efficiently (p-value = 0.0017, paired sample t-test) bringing 94.0±2.0% of reads mapping to rRNA down to 26.5±6.2% of reads mapping to rRNA, which represents a 13.0±4.1 fold enrichment of non-rRNA reads across the three strains (Fig. 3a, Supplementary Fig. 1). We also compared the performance of our method to the Ribo-Zero bacterial kit, which we applied to the same RNA samples. In 2 of 3 cases, our technique yielded higher performance than Ribo-Zero (10.0, 17.6 and 11.5-fold enrichment of non-rRNA reads by our RNase H method versus 3.6, 9.6 and 15.3-fold enrichment by Ribo-Zero, respectively). The overall efficiency was uniform across the three strains, highlighting the consistent performance of strain-specific probe sets. In addition, for all non-rRNA reads, depleted samples showed consistent overall mappability (79.2±12.5% for depleted samples, 75.8±8.1% for un-depleted samples, Supplementary Fig. 2) and a higher proportion of reads mapped to CDSs (93.6±1.4% for depleted samples, 83.8±2.9% for un-depleted samples, Supplementary Fig. 2), indicating our method minimally affects overall library quality. Together these results demonstrate that the RNase H based rRNA depletion method has favorable performance across three diverse strains.

**Figure 3:**
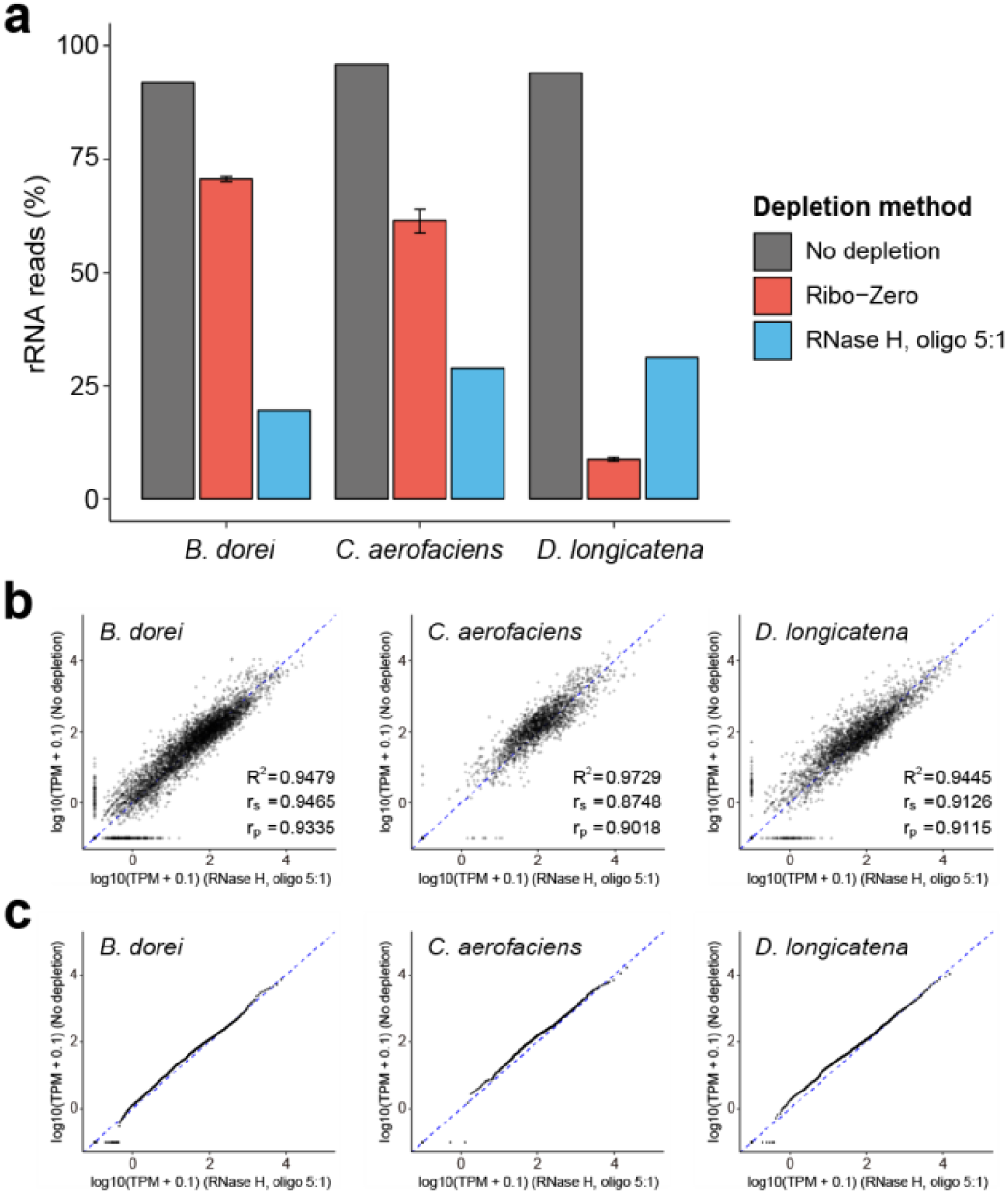
Application of RNase H based rRNA depletion with oligos to three diverse gut microbiota species. **a)** Proportion of rRNA-aligning reads of total mapped reads (% rRNA reads), for un-depleted, Ribo-Zero depleted, and RNase H depleted samples under optimized reaction condition across three microbiota species from different phyla. The same RNA samples used for Ribo-Zero depletion were pooled together to yield the RNA used for RNase H reaction. **b, c)** Consistency in transcriptome between rRNA depleted samples and un-depleted samples in terms of expression correlation **(b)** and expression distribution **(c)**. TPM indicates transcripts per million for each CDS. **c)** Points lie on a straight line in Quantile-Quantile (Q-Q) plots if there is no global shift in the distribution of expression profile between depleted samples and un-depleted samples.

Next, to determine if rRNA depletion may have altered the transcriptome due to off-target depletion, we compared the depleted expression values to un-depleted expression values for all annotated CDSs. Importantly, since both samples were generated from the same RNA sample, any difference reflects bias due to the depletion method (in addition to technical bias associated with sample preparation). We mapped at least 600K reads to annotated CDSs for each sample, to ensure high coverage for expression quantification (Supplementary Table 2). Overall, the depleted samples showed high consistency with un-depleted samples in terms of transcriptome correlation (R2 = 0.948, 0.973, 0.944, respectively, Fig. 3b and Supplementary Fig. 3) and expression distribution (quantile-quantile plot, points lie on a straight line if there is no global shift in expression profile) (Fig. 3c and Supplementary Fig. 4). Together, these results validate that the RNase H based rRNA depletion method minimally perturbs the underlying transcriptome.

### Amplicon-based probe generation and validation

The chemically synthesized oligo probe strategy provides efficient rRNA depletion but requires synthesis of a specific probe library. While the cost of the probe library can be amortized over many reactions and the total cost is minimal compared to purchasing a commercial kit (Table 1), for small scale studies it may be desirable to generate probes on a smaller scale for a lower overall cost. We therefore set out to validate a PCR amplicon and ssDNA generation strategy to generate probes from a microbial DNA template derived from universal primers. In theory, this strategy could also allow for the generation of complex probe sets from templates of multiple different species. Using universal primers where the forward primer is 5’-phosphorylated (Supplementary Table 3), we amplified the 16S and 23S regions from *B. dorei* and subjected the amplicons to lambda exonuclease digestion, which selectively digests the 5’-phosphorylated strands(46) to yield complementary ssDNA probes (amplicon) (Methods). We then repeated the testing of rRNA depletion efficiency on the same *B. dorei* RNA as in Fig. 3, utilizing the same reaction conditions and a 5:1 or a 10:1 amplicon probe-to-RNA ratio (Fig. 4a). The 5:1 ratio amplicon probes showed similar depletion efficiency (16.1% of reads mapping to rRNA) as the oligo probes, while the 10:1 ratio showed increased depletion efficiency (4.0% mapping to rRNA), which demonstrate that amplicon-based probes can also efficiently deplete rRNA. As before, we compared the expression values of depleted samples with the amplicon probes to expression values from the un-depleted samples generated from the same RNA (Fig. 4b-c). The amplicon-based depleted samples showed high consistency with un-depleted samples (R2 = 0.936). Together, these results demonstrate that in addition to chemically synthesized oligonucleotides, ssDNA probes generated from PCR amplicons can yield efficient rRNA depletion with minimal perturbation of the transcriptome.

**Figure 4:**
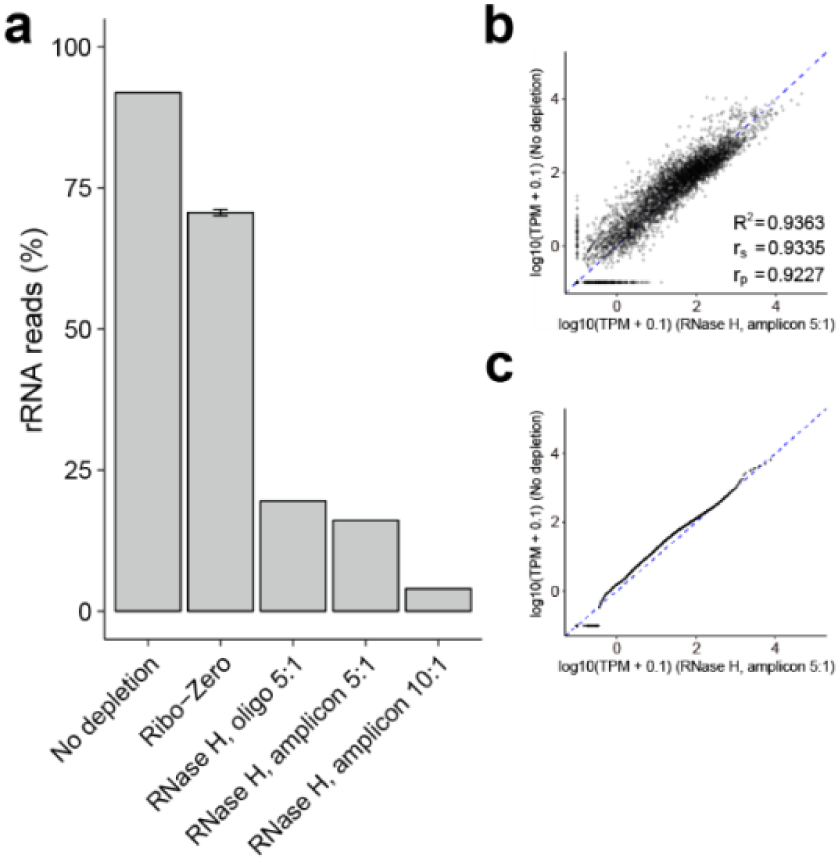
RNase H based rRNA depletion with amplicons. **a)** Proportion of rRNA-aligning reads of total mapped reads (% rRNA reads), for un-depleted, Ribo-Zero depleted, and RNase H depleted samples with chemically synthesized oligonucleotides (oligo) or amplicon-based ssDNA (amplicon) probes. **b, c)** Scatter plot **(b)** and Quantile-Quantile (Q-Q) plot **(c)** for depleted sample using amplicon probes with 5:1 probe-to-RNA ratio and un-depleted sample.

### Applicability of probes to closely related species

The cost of generating new probe libraries for novel species may impose practical experimental constraints for studies that involve transcriptomic profiling of many different microbial strains. We hypothesized that probe libraries developed for a specific strain may be applied also to closely related species with similar 16S and 23S sequences. A single probe library that can be scalably re-applied to many potential strains could thus significantly reduce the overall cost. We therefore tested the *B. dorei* oligo probe pool on two closely related *Bacteroides* species, *Bacteroides uniformis* ATCC 8492 and *Bacteroides vulgatus* ATCC 8482 with the same library preparation and optimal reaction conditions as before. In addition, we tested the same probe library on distantly related species, the *C. aerofaciens* and *D. longicatena* strains tested before. The *B. dorei* probe library was able to efficiently deplete *B. uniformis* and *B. vulgatus* rRNA (Fig. 5a) with minimal perturbation to their transcriptomes (R2 = 0.972, 0.971, respectively, Supplementary Fig. 3 and Supplementary Fig. 5) as before. As expected, the probes did not deplete rRNA from the distantly related organisms (*C. aerofaciens* and *D. longicatena*).

**Figure 5:**
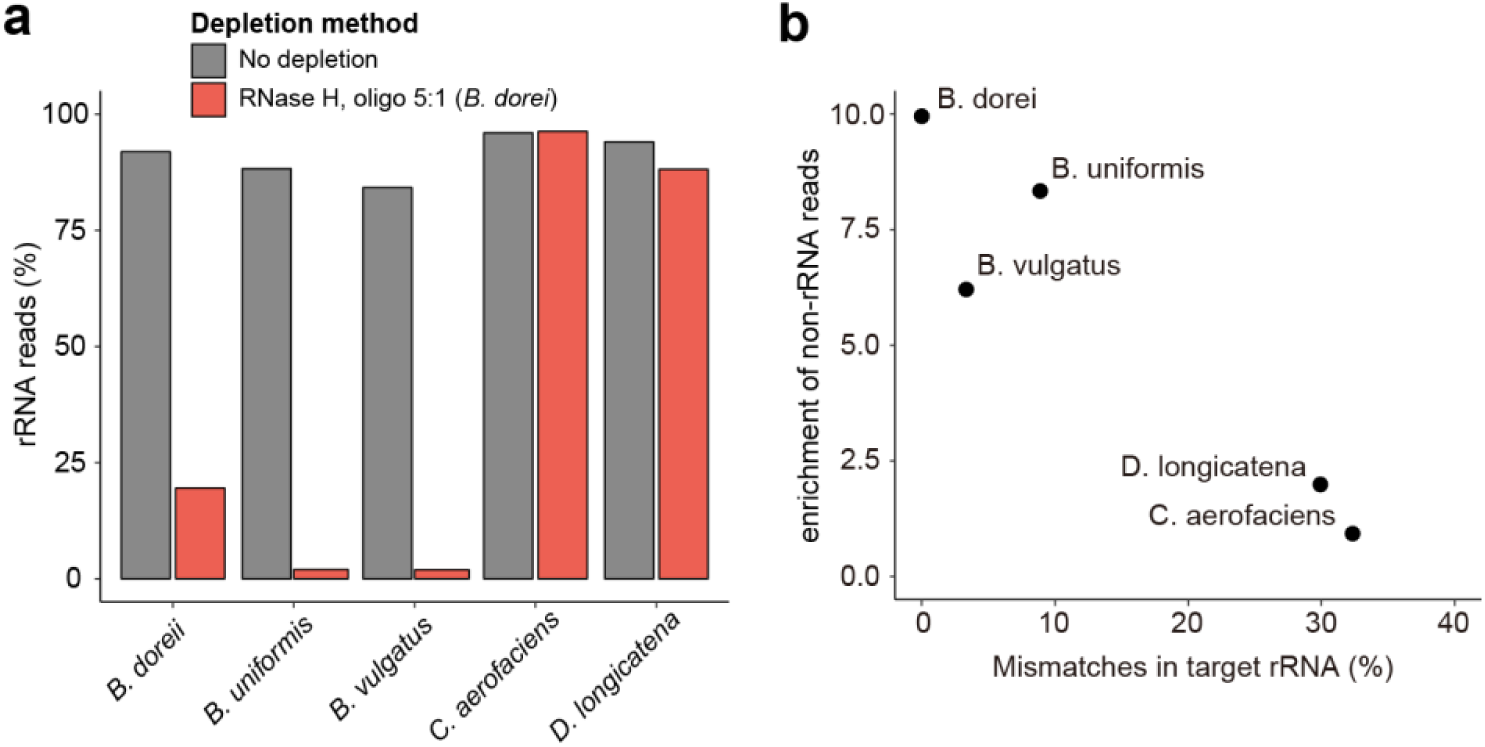
Oligo probes can be applied to closely related species. **a)** Proportion of rRNA-aligning reads of total mapped reads (% rRNA reads), for un-depleted and RNase H depleted samples using an oligo probe library designed for *B. dorei* across closely related species (*B. uniformis* and *B. vulgatus*) and distantly related species (*C. aerofaciens* and *D. longicatena*). **b)** Scatter plot between probe-to-target sequence similarity and fold enrichment of non-rRNA reads for five species. Probe-to-target sequence similarity was calculated as the average of percentages of base with mismatches in 16S and 23S rRNA alignments.

To understand the correlation between probe-to-target sequence similarity and rRNA depletion efficiency, we quantified the percent of mismatches in target rRNA sequences compared to *B. dorei* rRNA sequences. Species with less than 10% mismatches overall could be efficiently depleted (Fig. 5b). We performed a similar analysis for each individual oligo probe, and the correlation of probe mismatches to depletion efficiency of the cognate target sequence suggests that probes with less than 25% mismatches can be used to efficiently deplete local regions of rRNA (Supplementary Fig. 6). Taken together, these results demonstrate scalability of our designed probe libraries for reuse in closely related species.

### Application of rRNA depletion in non-model isolate transcriptome profiling

The RNase H-based rRNA depletion technique we have developed enables large-scale transcriptomic interrogation of non-model organisms. To illustrate this utility, we investigated dietary sugar utilization and microbial cross-feeding of a novel gut microbiota isolate, *B. dorei*. Short chain fatty acids (SCFAs) are considered primary end-products of carbohydrate fermentation by gut microbes(47) but mechanistic details around their production from various carbohydrate substrates and potential for cross-feeding has not been systematically delineated; transcriptional analysis offers a novel hypothesis-generating approach towards this goal. We thus selected 14 diverse carbohydrates (6 monosaccharides, 4 disaccharides and 4 pectins) and 5 SCFAs (Fig. 6a and Supplementary Table 7) to expose to the strain *in vitro* to characterize specific transcriptional responses towards this goal.

**Figure 6:**
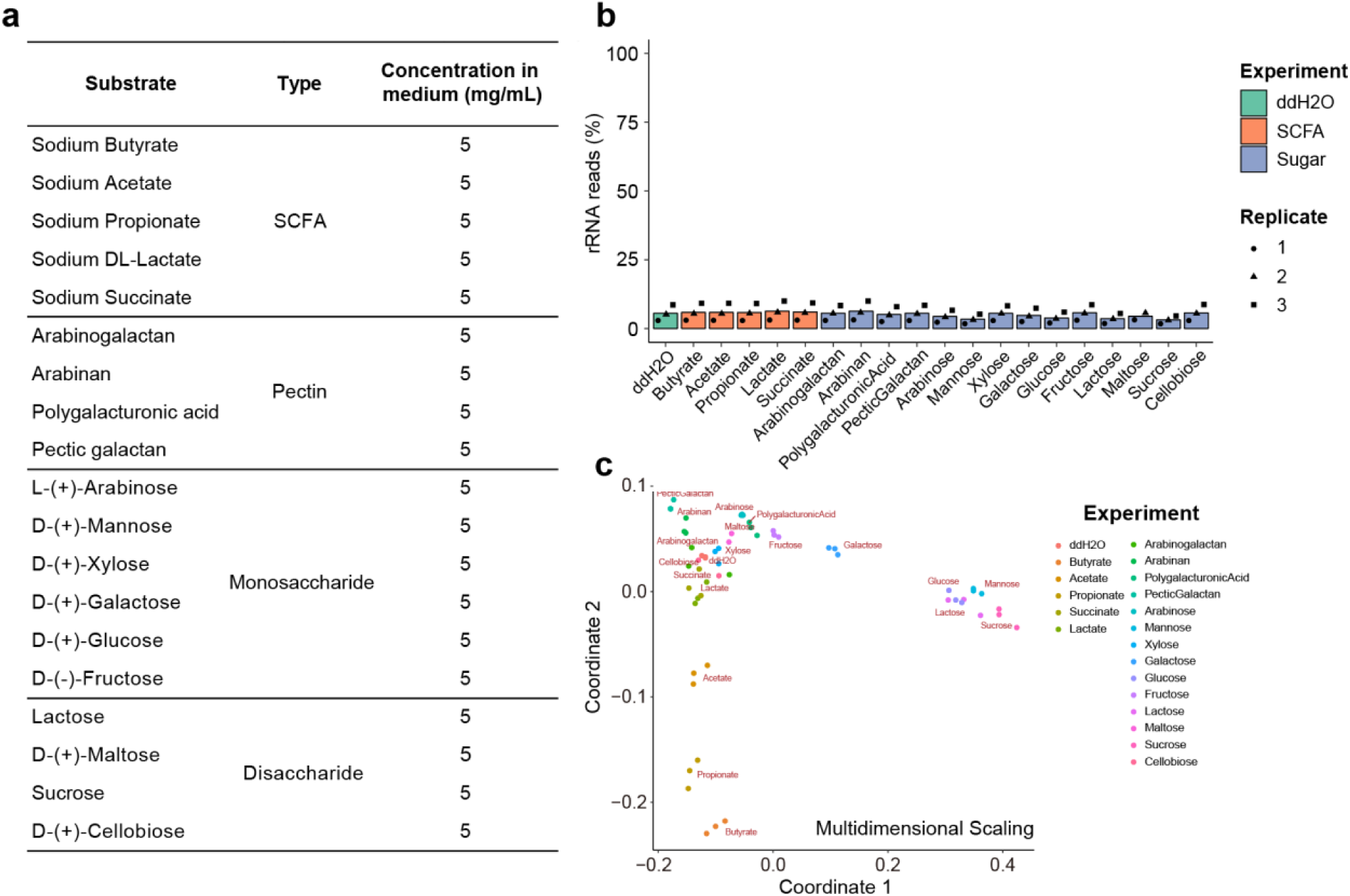
High-throughput microbial RNA-seq screening of *B. dorei* on different substrates. **a)** The 5 short-chain fatty acids (SCFAs) and 14 carbohydrates tested. *B. dorei* cultures were treated with each substrate at 5 mg/mL final concentration during exponential growth phase. **b)** The RNase H based method showed efficient and consistent rRNA depletion on pooled RNA samples from different conditions. Three biological replicates were performed for each condition. **c)** Multidimensional scaling ordination of 20 conditions based on overall transcriptomes. Pairwise Spearman correlations between conditions were calculated using expression profiles and multidimensional scaling was then performed on normalized Spearman correlations to visualize the impact of substrates on bacterial transcriptome.

*B. dorei* cultures at exponential phase were treated with these compounds for 90 minutes, and library preparation was performed as before; samples from different conditions were ligated to uniquely barcoded adapters following the standard RNAtag-seq protocol(23). RNase H-based rRNA depletion was then performed on pooled barcoded RNA utilizing the optimal reaction conditions and the *B. dorei* oligo probe library (Methods). As expected, our method yielded efficient and consistent rRNA depletion on pooled RNA samples from different conditions across replicates with 4.97±2.46% of reads mapping to rRNA (Fig. 6b). Moreover, biological replicates after depletion showed high consistency in transcriptome profiles (Supplementary Fig. 7), indicating the high reproducibility of our method.

Next, to evaluate the overall impact of each substrate on gene expression of *B. dorei*, transcriptome profiles of different conditions were subjected to ordination by multidimensional scaling (Fig. 6c). Interestingly, this analysis revealed significant transcriptome perturbation by glucose and mannose as well as two glucose-containing disaccharides (lactose and sucrose). In addition, the SCFAs acetate, propionate and butyrate (but not lactate and succinate) resulted in large-scale transcriptional shifts. In-depth analyses of specific differentially expressed genes and polysaccharide-utilization loci (PULs) revealed compound-specific transcriptional responses and pathways which likely mechanistically underlie the organism’s ability to metabolically utilize these compounds (Supplementary Fig. 8, Supplementary Table 8 and Supplementary Table 9). Importantly these results illustrate the ability of large-scale transcriptomics (enabled by our RNase H depletion approach) to characterize the metabolic and transcriptional regulatory capacity of non-model organisms in a rapid and cost-effective manner.

## Discussion

To facilitate use of this rRNA depletion technique by the research community, we have included a detailed step-by-step protocol for the method (Supplementary Methods). In addition, to streamline design of oligonucleotide pools, we have created a simple tool to generate probe libraries from target 16S and 23S sequences (https://github.com/hym0405/RNaseH_depletion). The tool can also perform simple calculations of probe identity to various different 16S and 23S sequences to evaluate the ability of pools to be applied to different sequences, and potential off-target sites prediction based on sequence similarity. For all five bacterial species we performed rRNA depletion on, we used this tool to evaluate potential off-target sites (Methods). We observed no off-target transcripts for four strains, with only one transcript (JDJECPLG_03071) identified as off-target for *D. longicatena*. This transcript is located in a fragmented contig of 1172bp and is perfectly identical to a part of *D. longicatena* 23S rRNA sequence, and is therefore likely a truncated rRNA gene that was not successfully annotated. Moreover, probe sets designed by the tool for common microbes (5,467 representative bacterial genomes from the RefSeq database) are also provided (Supplementary Table 10).

We developed two separate strategies for generating probe libraries that perform with similar efficiency. While chemically synthesizing probes requires less hand-on time, it constitutes a high up-front cost (~$450/probe set) but generally yields enough material for a large number of reactions (>12,000 reactions). Alternatively, in-house amplicon-based ssDNA probes do not require a significant up-front cost, but do require more time (~8-hour protocol end-to-end) to generate and quality control. Based purely on cost (Supplementary Fig. 9), synthesized probes become cost-effective after ~90 reactions compared to amplicon probes. Importantly however, we note that the cost per reaction in both cases ($4.98 for amplicon probes, ~$0.04 for oligo probes and $12.90 for other reagents per depletion reaction, Supplementary Table 11) is significantly lower than commercially available kits (e.g. ~$80 per reaction for the discontinued Ribo-Zero rRNA Removal Kits, Table 1).

Beyond the application to single-species bacterial rRNA depletion shown here, the method may have utility for depletion of unwanted sequences for various transcriptomics applications. For example, our approach could be applied to non-model organisms for rRNA depletion. In addition, extension of this strategy to more complex probe pools could enable high efficiency rRNA depletion for metatranscriptomics experiments. Chemically synthesized oligo pools could be designed for specific environments based on the expected genera present, or custom probe pools could be generated on a per-sample basis by pooled amplification of 16S and 23S rRNA from a complex microbial community.

Finally, we note that for specific applications, further optimization to reaction times, probe-to-RNA ratios, and RNase H reaction temperature could even further improve efficiency. We chose to use conservative reaction conditions that still achieved significant depletion of rRNA and enrichment of non-rRNA reads (Supplementary Fig. 10) to avoid any potential off-target effects. However, future work could further validate and improve upon different reaction conditions in the context of specific samples to yield accurate transcriptome data.

We have developed a new RNase H based ribosomal RNA depletion method for bacterial transcriptomics that is compatible with barcoded or fragmented RNA libraries. We validated that the method has high efficiency for rRNA depletion (~13-fold enrichment of non-rRNA reads) and minimal off-target perturbation to the transcriptome on three diverse bacterial species. We showed that the method is compatible with chemically synthesized oligo probes or amplicon-based ssDNA probes generated from genomic DNA. Finally, we demonstrated that probe pools can be successfully applied to closely related species. This technique is expected to have broad utility in routine bacterial transcriptomics experiments and emerging large-scale bacterial RNA-seq studies.

## Data Availability

Code utilized in this study can be accessed at https://github.com/hym0405/RNaseH_depletion.

The sequencing data generated in this study have been submitted to the NCBI BioProject database (http://www.ncbi.nlm.nih.gov/bioproject/) under accession number PRJNA542677.

## Funding

This work was supported by National Institutes of Health to H.H.W. [1R01AI132403, 1R01DK118044]; Burroughs Welcome Fund to H.H.W. [PATH 1016691]; Bill & Melinda Gates Foundation to H.H.W. [INV-000609]; the Schaefer Research Scholars Program to H.H.W.; a Fannie and John Hertz Foundation Fellowship to R.U.S.; and a NSF Graduate Research Fellowship to R.U.S. [DGE-1644869].

## Author Contributions

Y.H. and R.U.S. designed the method, performed experiments and analyzed all data. A.K. assisted with experiments. H.H.W. supervised the project. All authors wrote and approved the manuscript.

## Disclosure Declaration

H.H.W., Y.H. and R.U.S. are inventors on a provisional patent application filed by the Trustees of Columbia University in the City of New York regarding this work. H.H.W. is a member of the Scientific Advisory Board of SNIPR Biome.

## Acknowledgements

We thank Nathan Johns and Peter Sims for helpful discussions.

## Supplementary Table captions

**Supplementary Table 1:**

Sequences of oligonucleotide probes for *B. dorei*, *C. aerofaciens* and *D. longicatena*

**Supplementary Table 2:**

Coverage and correlation of transcriptome

**Supplementary Table 3:**

Primer sequences used for ssDNA generation

**Supplementary Table 4:**

Sequences used for adapter trimming

**Supplementary Table 5:**

Metadata for sequencing data

**Supplementary Table 6:**

Primer sequences used for library preparation

**Supplementary Table 7:**

Carbohydrates and SCFAs used in *B. dorei* screening experiment

**Supplementary Table 8:**

Annotation of up-regulated genes in *B. dorei* for SCFAs

**Supplementary Table 9:**

Annotation of susC/D gene pairs and PULs in *B. dorei*

**Supplementary Table 10:**

Probe sets designed for 5467 representative bacterial genomes in RefSeq database

**Supplementary Table 11:**

Cost analysis for RNase H based rRNA depletion method

