## Supplementary Methods for "Scalable and cost-effective ribonuclease-based rRNA depletion for transcriptomics"

**Amplicon ssDNA probe generation:**

Notes: To generate amplicon-based probes, we used bacterial ribosomal DNA (rDNA) universal primers from RNA-seq enrichment protocol of Human Microbiome Project(1).

16S amplicon: 5' phosphorylated forward primer 8F and reverse primer 1541R

23S amplicon: 5' phosphorylated forward primer 10F and reverse primer 2756R

Primer Sequence (5' to 3'):

16S_8F: /5Phos/ AGAGTTTGATCCTGGCTCAG

16S_1541R: AAGGAGGTGATCCAGCCGCA

23S_10F: /5Phos/ YGGTGGATGCCTTGGC

23S_2756R: YRCTTAGATGCTTTCAGCRBTTATC

1. First-round PCR (gDNA as template):
    a. 1X 50μL PCR reaction for both 16S and 23S
    b. Reaction setup (50μL):
        i. 1μL genomic DNA template
        ii. 2.5μL 5'-phosphorylated forward universal primer (8F for 16S gene, 10F for 23S gene)
        iii. 2.5μL reverse universal primer (1541R for 16S gene, 2756R for 23S gene)
        iv. 19μL nuclease-free water [ThermoFisher AM9937]
        v. 25μL Q5® Hot Start High-Fidelity 2X Master Mix [NEB M0494L]
    c. Thermocycling condition:

| Step | Temp | Time |
|---|---|---|
| Initial denaturation | 98°C | 30 seconds |
| 30 cycles | 98°C | 10 seconds |
| | 65°C for 16S | 20 seconds |
| | 61°C for 23S | |
| | 72°C | 2 minutes |
| Final extension | 72°C | 2 minutes |
| Hold | 4°C | ∞ |

    d. Gel extraction:
        i. Perform gel electrophoresis (E-Gel™ EX Agarose Gels, 2% [ThermoFisher G402002])
        ii. Excise DNA bands (~1500bp for 16S and ~2800bp for 23S) and gel extract with the Wizard™ SV Gel and PCR Cleanup System [Promega A9282] following the manufacturer's instructions.

2. Second-round PCR (gel extraction as template):
    a. 10x 50μL PCR reactions for both 16S and 23S
    b. Reaction setup (50μL):
        i. 1μL gel extraction product
        ii. 2.5μL 5'-phosphorylated forward universal primer (8F for 16S gene, 10F for 23S gene)
        iii. 2.5μL reverse universal primer (1541R for 16S gene, 2756R for 23S gene)
        iv. 19μL nuclease-free water
        v. 25μL Q5® Hot Start High-Fidelity 2X Master Mix
    c. Thermocycling condition:

| **Step** | **Temp** | **Time** |
|---|---|---|
| Initial denaturation | 98°C | 30 seconds |
| 32 cycles | 98°C | 10 seconds |
| | 65°C for 16S | 20 seconds |
| | 61°C for 23S | |
| | 72°C | 2 minutes |
| Final extension | 72°C | 2 minutes |
| Hold | 4°C | ∞ |

    d. 1X SPRI beads cleanup
        i. Perform cleanup of PCR products with SeraPure SPRI beads with 1X ratio of SPRI beads to volume of PCR product(2,3). Cleanup is as per usual AmpureXP instructions with the following minor modifications: 80% ethanol rather than 70% ethanol, and the same 55μL nuclease-free water should be used to elude purified DNA from beads for 10 PCR reactions, resulting in ~50μL purified PCR product for both 16S and 23S.
        ii. Measure concentrations of purified PCR products using NanoDrop 2000c spectrophotometers (expected ~1000ng/μL for both 16S and 23S)

3. Lambda exonuclease digestion
    a. 5x 80μL digestion reactions for both 16S and 23S
    b. 5U of lambda exonuclease per 1000ng of DNA input (10U/μL stock)
    c. Reaction setup (80μL):
        i. 10μL purified PCR product (~10μg DNA)
        ii. 8μL 10X Reaction Buffer [ThermoFisher EN0562]

        iii. 5μL(50U) lambda exonuclease [ThermoFisher EN0562]
        iv. 57μL nuclease-free water
    d. Incubate at 37 °C for 30 minutes, then stop reaction by heating at 80 °C for 10 minutes
    e. 1X SPRI beads cleanup
        i. Perform cleanup of PCR products with SeraPure SPRI beads with 1X ratio of SPRI beads to volume of digestion product(2,3). Cleanup is as per usual AmpureXP instructions with the following minor modifications: 80% ethanol rather than 70% ethanol, and the same 22μL nuclease-free water should be used to elude purified ssDNA from beads for 5 reactions, resulting in ~20μL purified amplicon ssDNA for both 16S and 23S.
        ii. Measure concentrations of amplicon ssDNA using Qubit™ ssDNA Assay Kit [ThermoFisher Q10212] (expected ~1500ng/μL for both 16S and 23S)

4. Mix 23S amplicon ssDNA and 16S amplicon ssDNA in equimolar amounts to generate ready-to-use amplicon probe mixes.
    a. **23S and 16S amplicon probe should be mixed equimolarly, e.g. 650ng 23S probe + 350ng 16S probe = 1000ng amplicon probe mix**

**RNase H based rRNA depletion:**

1. rRNA depletion (500ng total RNA as input)
    a. 15μL reaction setup:
        i. 500ng RNA
        ii. 2500ng ssDNA probe mix (chemically synthesized oligos or amplicon ssDNA)
        iii. 0.6μL 5M NaCl (200mM NaCl) [Fisher BP3581]
        iv. 1.5μL 1M Tris-HCl (100mM Tris-HCl pH 7.5) [ThermoFisher 15567-027]
        v. Add nuclease-free water to 15μL
        **vi. The probe-to-RNA ratio can be optimized if necessary. Generally, 5X probe is sufficient for efficient rRNA depletion.**
    b. Thermocycling condition (lid temperature: 105°C):
        **i. This step is important for probe annealing and will take approximately ~16 minutes to complete.**

| Temp | Time |
|---|---|
| 95°C | 2 minutes |
| -0.1°C/sec to 45°C | ~9 minutes |
| 45°C | 5 minutes |

    c. Spin down and immediately proceed to the next step.
    d. Prepare 5μL RNase H master mix:
        i. 3μL Hybridase™ Thermostable RNase H [Lucigen H39500]
        ii. 0.5μL 1M Tris-HCl (0.5μmol Tris-HCl pH 7.5)
        iii. 0.2μL 5M NaCl (1μmol NaCl)
        iv. 0.4μL 1M $MgCl_2$ (0.4μmol $MgCl_2$) [Sigma-Aldrich 208337]
        v. 0.9μL nuclease-free water
        **vi. The amount of RNase H enzyme can be optimized if necessary.**
    e. Mix RNase H master mix by pipetting up and down at least 10 times, spin down and preheat mix to 45°C, **use immediately**.
    f. add 5μL of preheated master mix to the 15μL reaction and mix by pipetting up and down at least 10 times.
    g. Spin down briefly and **immediately proceed to the next step**.
    h. Place samples in a thermocycler and incubate at 45°C for 30 minutes. (lid temperature: 60°C). After incubation, spin down the samples and place on ice. Proceed to the next step immediately.
        **i. The incubation time and temperature could be further optimized if necessary.**

2. RNA purification
   **Important Note:**
   - For RNA **not ligated with 3’ DNA adapter** (fragmented or unfragmented), probe removal by **DNase I digestion** is recommended;
   - For RNA **ligated with 3’ DNA adapter** (e.g. barcoded RNA in RNAtag-seq(4)), probe removal by **2X SPRI beads cleanup** is recommended.

   Probe removal by **DNase I digestion:**
   a. Prepare 30μL DNase I master mix, use immediately.
      i. 3μL TURBO™ DNase [ThermoFisher AM2239]
      ii. 5μL 10X TURBO™ DNase Buffer [ThermoFisher AM2239]
      iii. 22μL nuclease-free water
   b. Add 30μL DNase I master mix to 20μL RNase H reaction, mix by pipetting up and down at least 10 times.
   c. Place samples in a thermocycler and incubate at 37°C for 30 minutes. (lid temperature: 45°C). After incubation, spin down the samples and place on ice. Proceed to the next step immediately.
   d. Perform cleanup of DNase I digestion product with SeraPure SPRI beads with 2X ratio of SPRI beads to volume of depletion product(2,3). Cleanup is as per usual AmpureXP instructions with the following minor modifications: 80% ethanol rather than 70% ethanol, elution into 15μL nuclease-free water and removal of 14μL.

   Probe removal by **2X SPRI beads cleanup:**
   b. Perform cleanup of RNase H digestion product with SeraPure SPRI beads with 2X ratio of SPRI beads to volume of depletion product(2,3). Cleanup is as per usual AmpureXP instructions with the following minor modifications: 80% ethanol rather than 70% ethanol, elution into 26μL nuclease-free water and removal of 25μL.
   c. Perform cleanup of the product **again** with SeraPure SPRI beads with 2X ratio of SPRI beads to volume of the product(2,3). Cleanup is as per usual AmpureXP instructions with the following minor modifications: 80% ethanol rather than 70% ethanol, elution into 15μL nuclease-free water and removal of 14μL.

3. RNA samples after rRNA depletion can be further processed via standard microbial RNA-seq library preparation (e.g. in our paper, RNAtag-seq(4)) or utilized for other downstream applications. Alternatively, add 1uL SUPERase-In™ RNase Inhibitor [ThermoFisher AM2694] to each sample and store at -80°C.
