## Supplementary Figure for "Scalable and cost-effective ribonuclease-based rRNA depletion for transcriptomics"

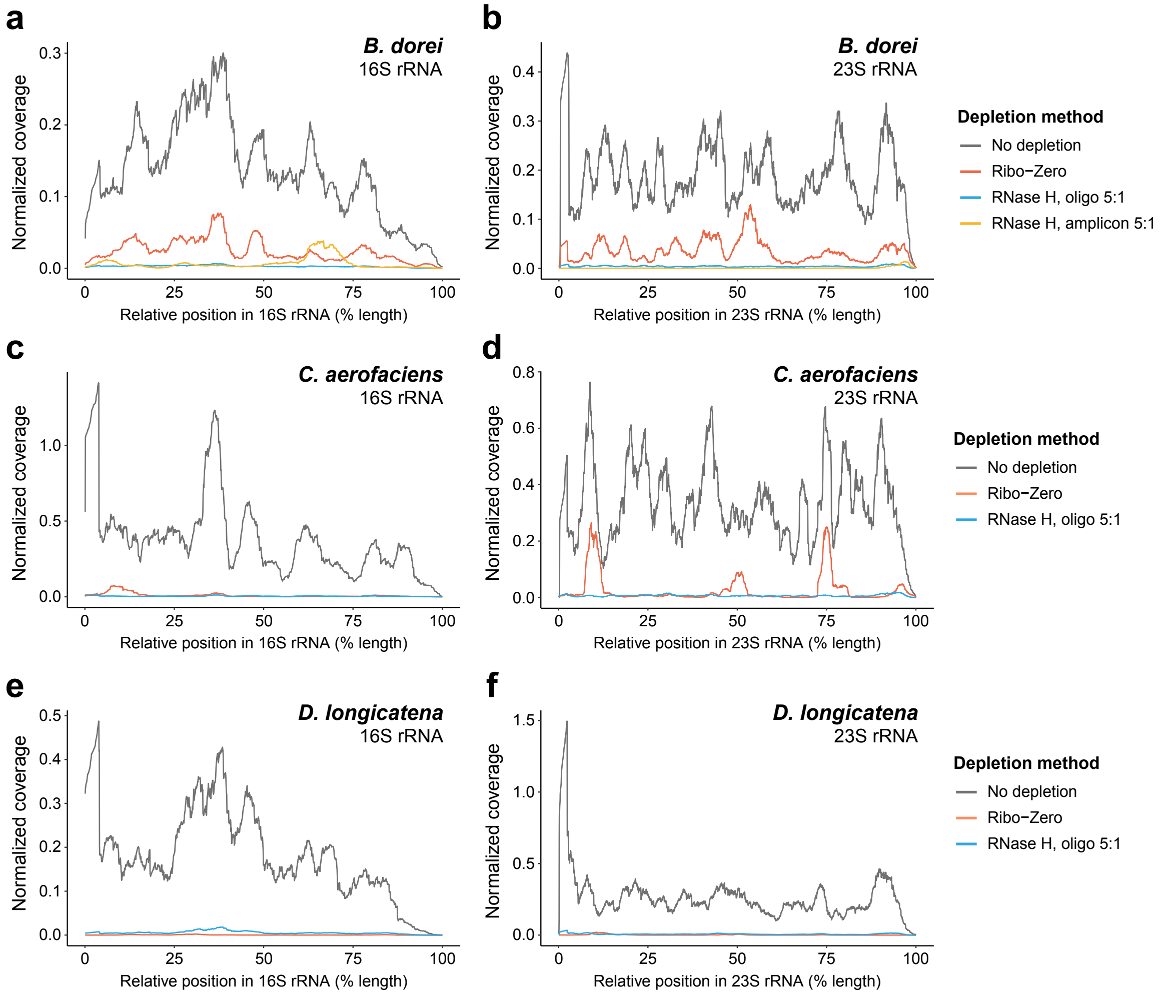


**Supplementary Figure 1: rRNA read coverage for three diverse gut species. a-f)** Normalized reads coverage along 16S and 23S rRNA for un-depleted, Ribo-Zero depleted and RNase H depleted samples of *B. dorei* **(a, b)**, *C. aerofaciens* **(c, d)** and *D. longicatena* **(e, f)**. RNase H depletion was performed with the optimized reaction conditions using oligo probe libraries and read coverage of each base was normalized to the number of non-rRNA mapped reads to avoid bias caused by the decreased proportion of rRNA-aligning reads.

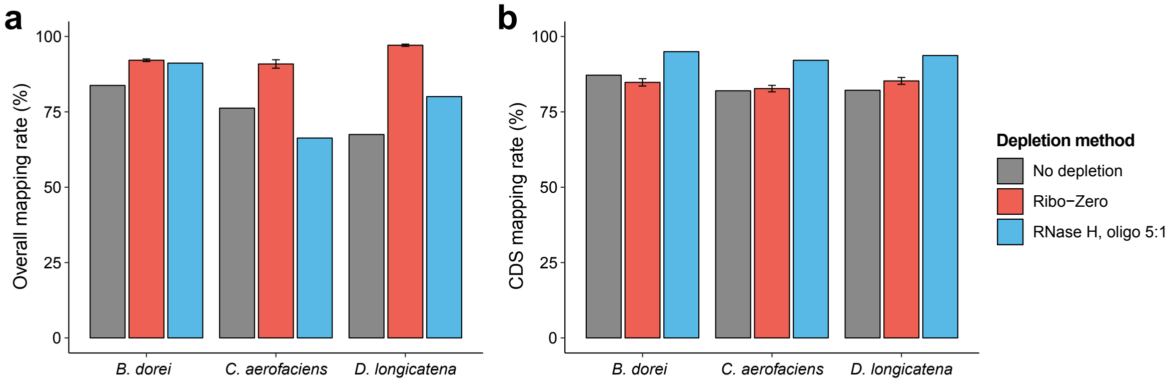


**Supplementary Figure 2: Metrics for non-rRNA read alignment. a, b)** Overall mapping rate of non-rRNA reads **(a)** and proportion of non-rRNA reads mapped to annotated CDSs **(b)** for un-depleted, Ribo-Zero depleted and RNase H depleted samples across the three species. RNase H depletion was performed with the optimized reaction condition using oligo probe libraries and all CDSs were annotated using Prokka with default settings.

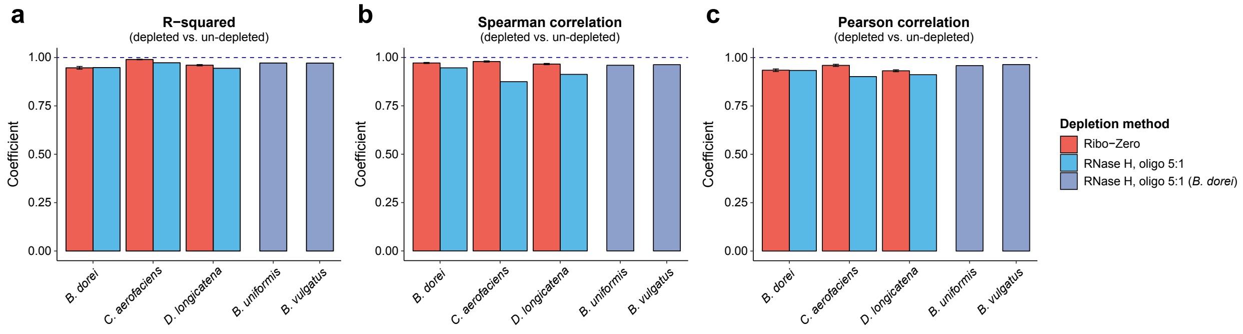


**Supplementary Figure 3. RNase H based rRNA depletion minimally perturbed underlying transcriptome based on three criteria of transcriptome consistency.** R-squared, Spearman correlation and Pearson correlation between rRNA depleted and non-depleted samples were calculated for different species and depletion strategies. For *B. uniformis* and *B. vulgatus*, rRNA was depleted using an oligo probe library designed for *B. dorei*.

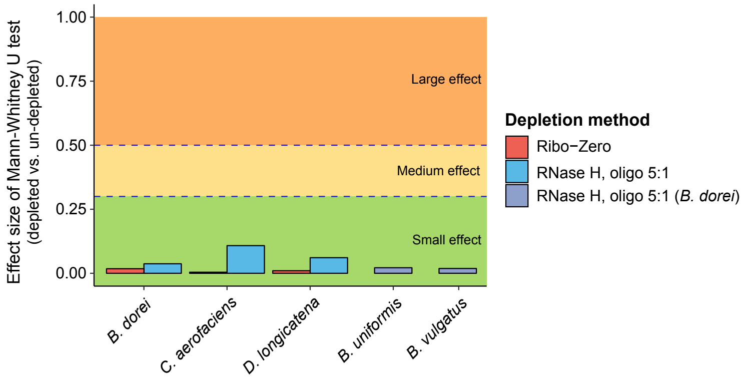


**Supplementary Figure 4. RNase H based rRNA depletion minimally altered the expression distribution of transcriptome.** Effect size *r* of Mann-Whitney U test was calculated between depleted and un-depleted samples to quantitively measure the distribution shift. For *B. uniformis* and *B. vulgatus*, rRNA was depleted using oligo probe library designed for *B. dorei*.

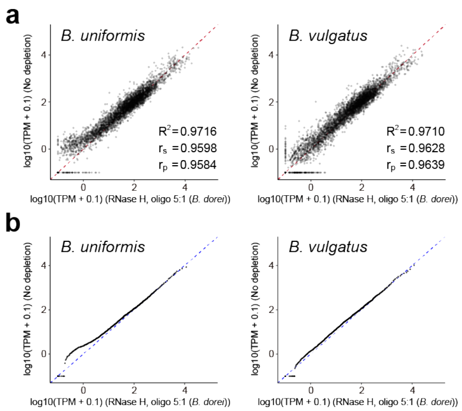


**Supplementary Figure 5: Consistency in transcriptome for two *Bacteroides* species after depletion using the *B. dorei* probe library. a, b)** Consistency in transcriptome between rRNA depleted samples and un-depleted samples in terms of expression correlation **(a)** and expression distribution **(b)** for *B. uniformis* and *B. vulgatus*. RNase H depletion was performed with the optimized reaction conditions using the *B. dorei* oligo probe library. TPM indicates transcripts per million for each CDS.

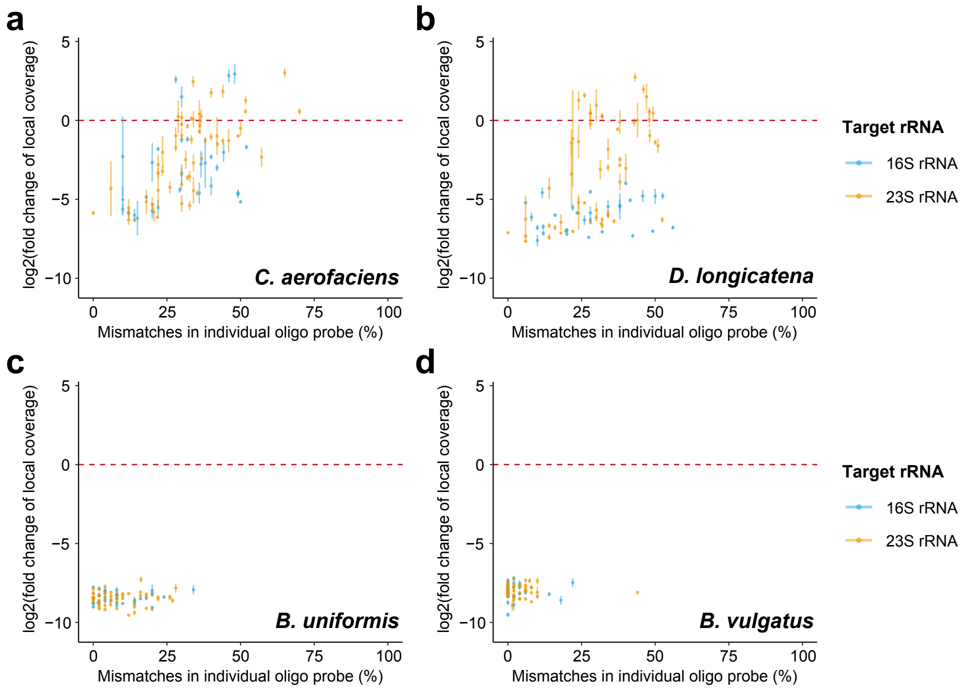


**Supplementary Figure 6: Correlation between probe mismatches and depletion efficiency for local sequences depleted with the *B. dorei* probe library. a-d)** Scatter plot between probe-to-target mismatches and depletion efficiency of the corresponding local rRNA sequence for two distantly related species, *C. aerofaciens* **(a)** and *D. longicatena* **(b)**, and closely related species, *B. uniformis* **(c)** and *B. vulgatus* **(d)**. Depletion efficiency of local rRNA sequence, i.e. fold change of local coverage, was calculated as the average fold changes of normalized coverage for each base in the region (reads coverage of each base was normalized to number of non-rRNA mapped reads).

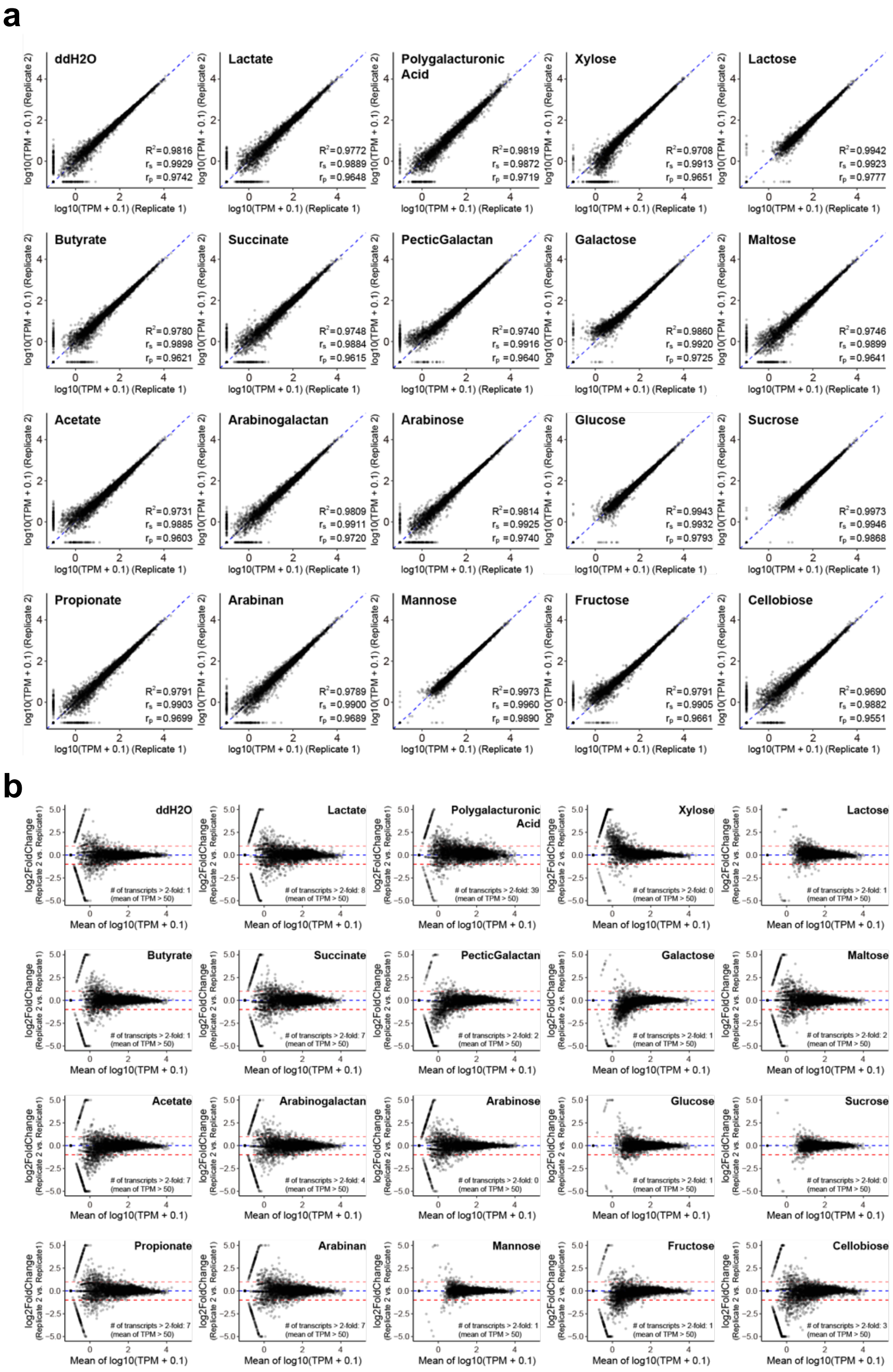


**Supplementary Figure 7. RNase H based rRNA depletion yielded high reproducibility between biological replicates. a, b)** Scatter plots **(a)** and MA plots **(b)** show consistent transcriptome profiles between biological replicates of *B. dorei* with the RNase H based depletion method. Numbers of relatively high-abundance transcripts (mean of TPM > 50) showing a fold change of TPM greater than 2 between two biological replicates were calculated for each condition in MA plots. TPM indicates transcripts per million for each CDS.

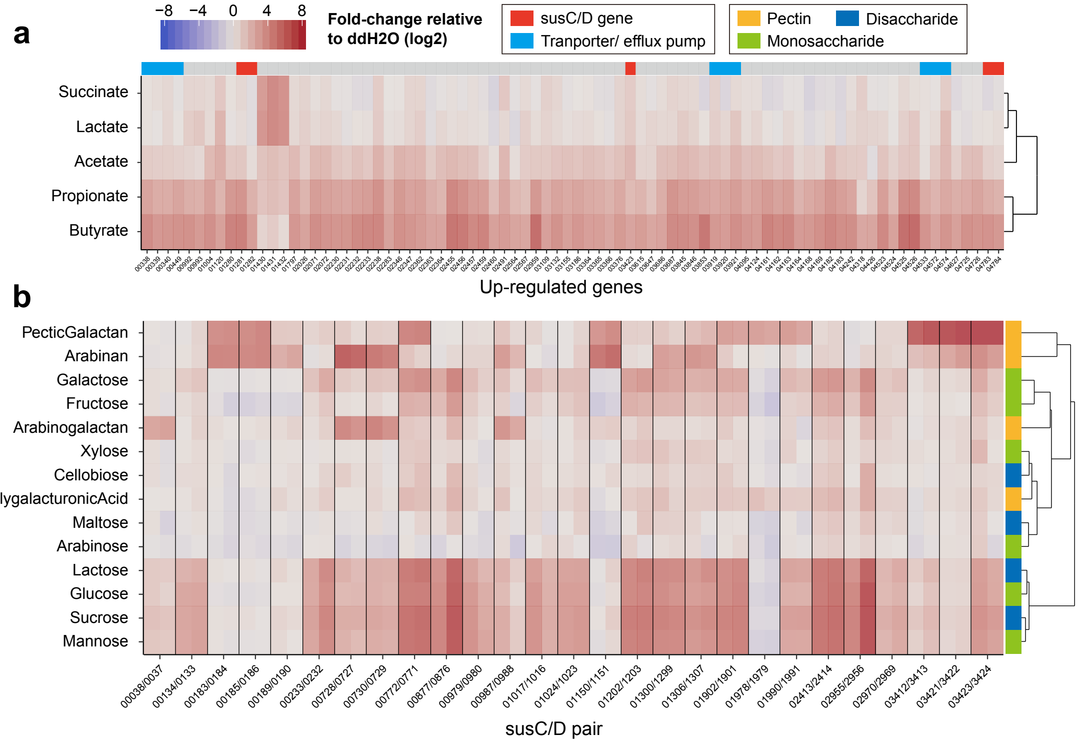


**Supplementary Figure 8: Heatmap of up-regulated genes in RNA-seq screening of *B. dorei* on different substrates. a)** Heatmap of fold-changes in expression level relative to ddH2O control for all 82 highly up-regulated genes (fold-change > 10 and adjusted p-value < 0.01 in at least one condition) across 5 SCFAs (for a detailed annotation of up-regulated genes, see Supplementary Table 8). **b)** Heatmap of fold-changes in expression level relative to ddH2O control for highly up-regulated 27 susC/D gene pairs identified in the *B. dorei* genome (fold-change > 10 and adjusted p-value < 0.01 in at least one condition) across 14 carbohydrates (for a full list of susC/D gene pairs and polysaccharide utilization loci prediction, see Supplementary Table 9).

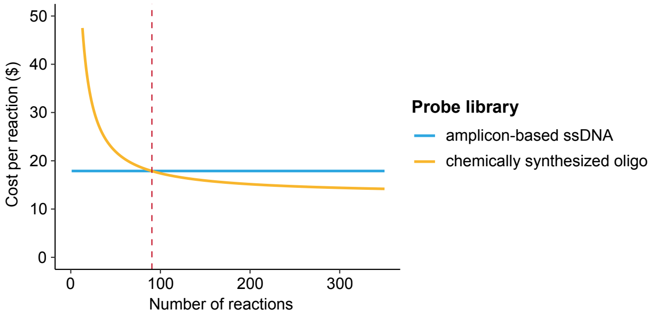


**Supplementary Figure 9: Cost per reaction for RNase H based rRNA depletion method.** Curve of estimated cost per reaction for RNase H based rRNA depletion using amplicon-based ssDNA probes ($17.88 per reaction) and chemically synthesized oligo probes ($12.90 per reaction for depletion reagents and $450 for up-front oligo probes synthesis). Chemically synthesized oligo probes become more cost-effective after ~90 reactions compared to amplicon-based ssDNA probes. This does not account for hands-on experimental time which should also be taken into account.

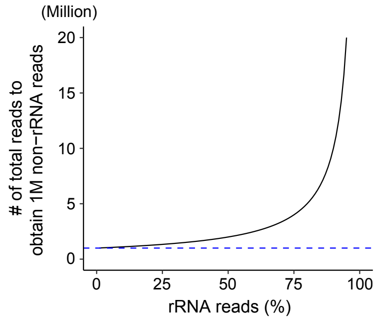


**Supplementary Figure 10. The theoretical relation between proportion of rRNA reads and the number of total reads required to obtain 1 million non-rRNA reads.** To obtain a certain sequencing depth of transcriptome, i.e. 1 million non-rRNA reads, the number of total reads required is inversely proportional to (1 – proportion of rRNA reads).
